# Paradigm shift for *cry* gene expression in *Bacillus thuringiensis*

**DOI:** 10.1101/2025.06.13.659657

**Authors:** Emilie Verplaetse, Leyla Slamti, Didier Lereclus

## Abstract

In most *Bacillus thuringiensi*s strains, the insecticidal *cry* genes are transcribed during sporulation by RNA-polymerases containing sigma factors E or K, leading to the formation of an insecticidal crystal within the mother cell along spore development. Some strains, such as the *kurstaki* HD1, a parent of commercial strains, also release insecticidal proteins Cry1I and Vip3A in the extracellular medium. *vip3A* gene expression is activated by the transcriptional regulator VipR at the onset of the stationary phase. Here, we expanded the VipR regulon to all the insecticidal-encoding genes of strain HD1 by identifying the VipR-binding box and conducting transcription assays. Unexpectedly, a VipR box was located in the promoter region of a putative N-acetylmuramoyl-L-alanine amidase gene upstream *cry1Ac in strain kurstaki* HD73, closely related to the HD1 but devoid of *vipR*. We demonstrated that introducing a functional *vipR* in this strain leads to the expression of the *amidase-cry1Ac* operon, resulting in an early and increased production of Cry1Ac. Investigating this phenotype, we showed that Cry1Ac was produced in a VipR-dependent manner in an HD73 Δ*spo0A* mutant. Similarly, an HD1Δ*spo0A* strain forms insecticidal crystals and produces all the insecticidal proteins encoded in its genome, including *cry2Ab* previously considered unexpressed in strain HD1. Finally, a genomic analysis revealed the presence of putative VipR-binding sequences upstream from insecticidal protein-encoding genes in several biopesticidal strains. Overall, our results break the dogma on the regulation of *cry1A* and *cry2A* genes and provide evidence of sporulation-independent Cry toxin production in biopesticidal Bt strains.

**Importance:** *Bacillus thuringiensis* is a remarkably efficient entomopathogen due to its ability to produce various insecticidal proteins, such as Cry or Vip. This property has made it a highly effective biopesticide used worldwide. Our work breaks the dogma of *cry1* and *cry2* genes being regulated solely by sporulation-specific sigma factors and thus exclusively expressed during this process. Indeed, we demonstrated that the VipR regulator controls the transcription of all the insecticidal protein genes encoded by a strain closely related to that of commercial biopesticides and turns on their expression at the beginning of the stationary phase, leading to the production of insecticidal crystals independently of sporulation. By providing new knowledge on the regulation of insecticidal protein genes, these findings bring new insight for the genetic improvement of Bt strains used as commercial biopesticides.

## Introduction

*Bacillus thuringiensis* (Bt) is a bacterium famous for its ability to produce a spore and massive amount of insecticidal proteins that will form a parasporal crystal inclusion (1, 2). The pathogenic properties of the bacterium towards insects rely on the production of Cry and Cyt toxins that have the property to crystallize and on the production and release of Vip, Sip and a few Cry toxins unable to form a crystal (3, 4). The majority of *cry*, as well as the *cyt* and *vip3* genes, are located on plasmids, and many Bt strains carry various combinations of insecticidal protein genes (1).

Since the first studies in the 1980s-1990s, a close association between *cry* gene expression and the sporulation process has been evidenced. Indeed, studies of *cry1A*, *cry4A*, *cry4B*, *cry11A*, *cry18,* and *cyt1A* gene expression have demonstrated that transcription of the insecticidal protein-coding genes was initiated from two overlapping promoters located upstream from the *cry* open reading frame (5–9), while *cry2* and *cry15* genes were transcribed from one promoter (10, 11). It was demonstrated that the *cry* and *cyt* promoters were transcribed by an RNA polymerase that contains the sporulation-specific sigma factor E (SigE) or K (SigK), active sequentially during the sporulation, that allows a sustained production of toxins in the mother-cell compartment during this process (9, 12, 13). Thus, it is widely admitted that expression of the *cry* and *cyt* genes encoding insecticidal proteins active against lepidopteran and dipteran insects is exclusively sporulation-dependent (14). This is particularly true of commercially-used Bt strains such as *kurstaki* HD1 and *israelensis*.

Besides this established link between sporulation and *cry* gene expression, a few observations questioning this paradigm have been reported without any explanations. Notably, the transcription of *cry1A*, *cry1E*, *cry1I*, and *cry2A* started before the onset of sporulation, and a faint production of Cry1 and Cry2 proteins was detected in a time frame that precedes the activation of SigE and SigK, in Bt *kenyae* and *kurstaki* strains isolated in Mexico (15). While the *cry1Ac* gene of Bt *kurstaki* HD73 is a typical example of a SigE and SigK-dependent crystal gene, it has been shown that a weak transcription of *cry1Ac* occurred in sporulation mutants of the HD73 strain (16). In a very different way from the genes (*e.g.. cry1*, *cry2* and *cry4*) coding for toxins active on Lepidoptera and Diptera, other modes of expression have been demonstrated for Bt strains active on other insect orders. The expression of the Cry3*-*coding gene is controlled by SigA and is independent of the sporulation sigma factors (17–19). Consequently, this coleopteran-specific pesticidal protein is produced from the end of the vegetative growth throughout the stationary phase (2). Lastly, the atypical production of Cry proteins by the LM1212 and HD977 strains has been unraveled by identifying CpcR, a specific transcriptional regulator located on a large plasmid (20, 21). In the LM1212 strain, the regulator leads to the production of Cry proteins only in a subpopulation that will not form a spore in a division of labor strategy (22).

The regulation of the insecticidal proteins that do not crystallize and that are liberated in the growth medium, such as Vip3A (4) or Cry1I (24), has remained unknown until the identification of VipR, the transcriptional regulator required for the expression of *vip3A*, in the HD1 strain (25). This transcription factor belongs to the PCVR family, together with the *B. anthracis* AtxA and the *Streptococcus pyogenes* Mga (26). The putative DNA binding site of VipR, designated VipR box, is an imperfect palindromic sequence, present in the *vip3A* promoter and in the promoter of the *vipR* gene, which is autoregulated (25). The VipR box was also found in the promoter region of other transcription units on the pBMB299 plasmid of the HD1 strain. Among these are the *cry1I*, *cry2Ab,* and the three-gene operon harboring *cry2Aa* suggesting that VipR may also be involved in expressing these *cry* genes. The VipR box was also identified in the promoter region of two N-acetylmuramoyl-L-alanine amidase-encoding genes or pseudogenes (25).

This study overturns the dogma that Cry1 and Cry2 insecticidal proteins are strictly produced during sporulation. We demonstrate an early production of Cry proteins independent of the sporulation genetic network but VipR-dependent in Bt strains that produce a functional VipR regulator. In summary, our study shows that, upon entry into the stationary phase, VipR controls the expression of all the insecticidal protein genes in *kurstaki* strain HD1.

## Results

### VipR controls the expression of all the insecticidal protein genes of the Bt *kurstaki* HD1 strain

The VipR transcriptional regulator of the Bt *kurstaki* HD1 strain positively controls its expression and the expression of the *vip3A* gene, located on the same plasmid (25). A conserved 32-bp sequence designated VipR box, whose integrity was critical for the expression of *vipR* and *vip3A,* was identified in their promoter region. This sequence was also identified in the promoter of 5 other transcription units on the pBMB299 plasmid: upstream of *cry2Ab*, *cry1I*, a three-gene operon containing *cry2Aa* and two genes annotated as N-acetylmuramoyl-L-alanine amidases (abbreviated as *ami*), with one *ami* gene being interrupted by a transposon (25). The consensus sequence of the VipR box was described using the IUPAC nucleotide code for the variable positions, and we used this sequence to search for other genes of the HD1 strain putatively regulated by VipR. No VipR box was found in the chromosome of the HD1 strain, but two additional occurrences of the VipR box were identified on plasmids. One sequence was located upstream from an *ami* gene on the pBMB65 (*ami65*), whereas the other was located upstream from an *ami* gene on the pBMB95 (*ami95*) (Fig. 1A and S1). Similarly to the other VipR boxes, the conserved sequences are located 17 nucleotides upstream from a putative Sig A -10 box. These results suggested that VipR was involved in regulating the expression of 7 transcriptional units in the HD1 strain, all plasmid-encoded. In order to confirm that VipR was involved in the expression of these putative targets, we analyzed the transcriptional activity of DNA fragments located upstream from the coding sequence of *cry2Ab*, *cry1I*, the first gene of the *cry2Aa* operon and the *ami95* gene located on the pBMB95. We used a heterologous host that does not harbor *vipR*, the Bt *kurstaki* HD73^-^ strain, which is genetically close to the HD1 strain. HD73^-^ is cured of the *cry1Ac*-carrying pHT73 plasmid (Table 1). Therefore, it does not produce any Cry toxin (27). The DNA fragments were fused to the promoterless β-galactosidase *lacZ* gene present on the pHT304.18Z, abbreviated as pHT-P*_gene_* (Table 1) (28). Each plasmid was introduced in the HD73^-^ strains carrying the pP*_xyl_*-*vipR* plasmid, allowing the expression of *vipR* under the control of the xylose-inducible promoter of *xylA*, or carrying the empty pP*_xyl_* plasmid (Table 1) (25). The HD73^-^ (pP*_xyl_*-*vipR*, pHT-P*_gene_*) and the HD73^-^ (pP*_xyl_*, pHT-P*_gene_*) strains (Table 2) were patched on LB plates containing X-gal and xylose. Blue colonies were observed after 24 h of growth for all the strains that carry the pP*_xyl-_vipR*, while the colonies were white for the pP_xyl_-carrying strains, indicating that the bacteria produced β-galactosidase activity only when the regulator was produced (Fig. 1B). A light blue coloration was observed for the HD73^-^ (pP*_xyl_*, pHT-P*_cry2Aa_*) strain after a prolonged incubation probably as a result of the activation of the SigE-dependent promoter included in the DNA cloned upstream of *lacZ* (Fig. S2) (12). This demonstrates that VipR positively regulates the expression of the HD1 genes that contain a VipR box in their promoter.

**Fig. 1.**
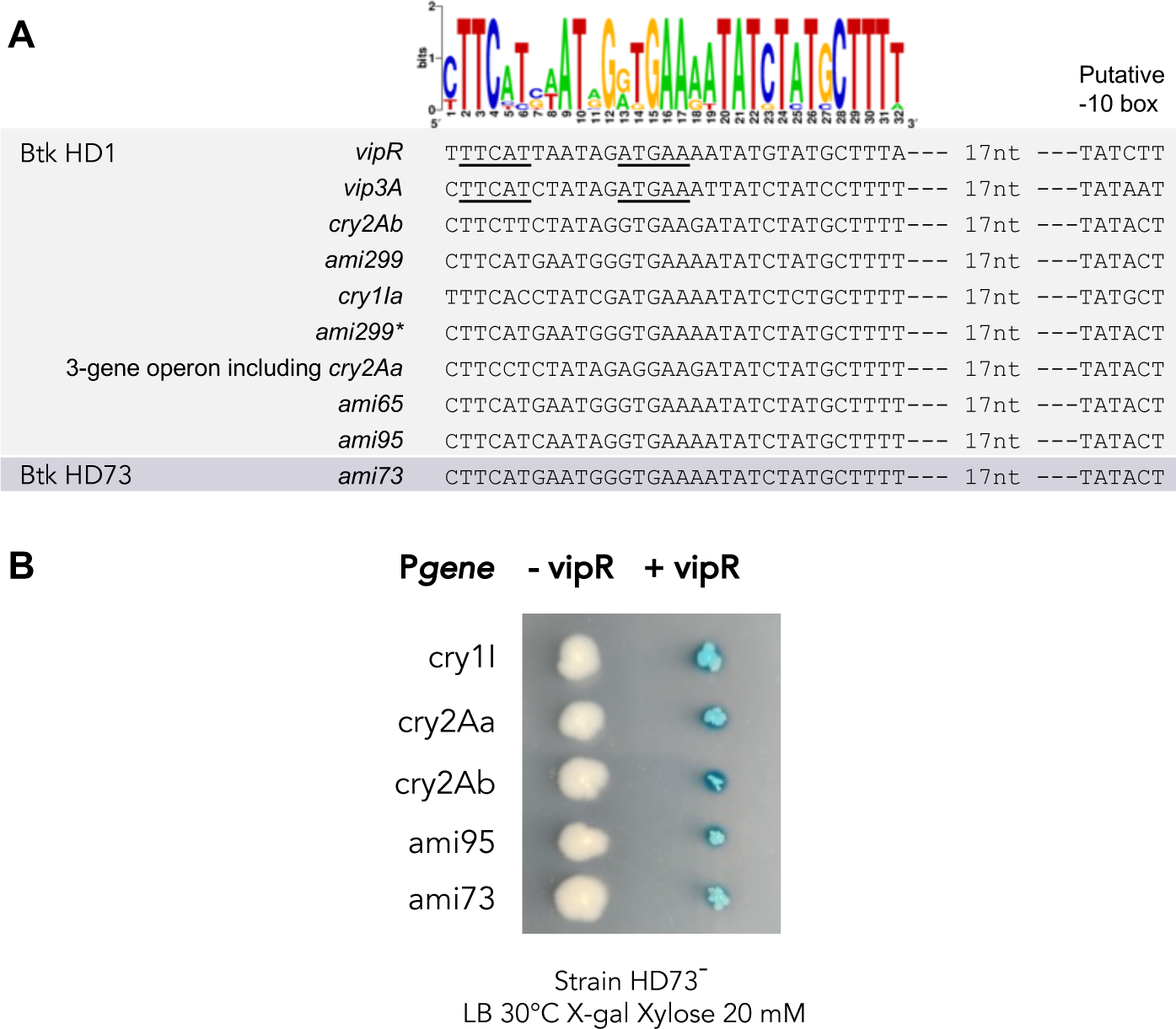
VipR controls the expression of the insecticidal protein genes containing a VipR box in their promoter. **A.** Alignment of the VipR boxes present on the pBMB299, pBMB65 and pBMB95 of the HD1 strain and on the pHT73 of the HD73. The name of the gene that contains the conserved sequence in its promoter is indicated on the left. The distance between the putative -10 box and the last nucleotide of the box is indicated. A consensus sequence is shown on top of the alignment. Figure adapted from (25). **B.** The strains HD73^-^ harboring the plasmids pP*_gene_*-*lacZ* and the pHT-P*_xyl_*-*vipR* (column « + vipR ») or the pHT-P*_xyl_* (column « - vipR ») were patched on LB medium containing X-gal (50 µg/mL) and xylose 20 mM and grown at 30°C for 24 h. The name of the gene whose promoter is fused to *lacZ* is indicated on the left.

**Table 1:** Plasmids used in this study.

| Name | Description |
| --- | --- |
| pBMB299 | Native plasmid of the HD1 strain, carries the *vipR*, *vip3A*, *cry1IA*, *cry2Aa*, *cry2Ab* and *cry1Aa* genes |
| pBMB95 | Native plasmid of the HD1 strain, carries the *cry1Ac* gene |
| pHT73 | Native plasmid of the HD73 strain, carries the *cry1Ac* gene |
| pHT304.18Z | *E. coli*/Bt shuttle vector conferring resistance to ampicillin in *E. coli* and to erythromycin in Bt. Designed to allow transcriptional fusion with *lacZ* (28) |
| pP<sub>xyl</sub> | The pHT16.18K plasmid that contains the *xylR* repressor gene and the promoter of the *xylA* gene to allow gene expression upon xylose induction (51) |
| pP<sub>xyl-vipR</sub> | Plasmid allowing the expression of *vipR* under the xylose inducible promoter of *xylA* (25) |
| pHT-P<sub>cry1I</sub> | A 381-bp DNA fragment containing the *cry1I* promoter was PCR amplified using genomic of the HD1 strain as the template and the primer pair Pcry1Ia-F/ Pcry1Ia-R. The DNA fragment was ligated between the *HindIII* and *BamHI* sites of the pHT304.18Z |
| pHT-P<sub>cry2Aa</sub> | A 2108-bp DNA fragment containing the promoter of the first gene of the *cry2Aa* operon was PCR amplified using genomic of the HD1 strain as the template and the primer pair Pcry2Aa-F/ Pcry2Aa-R. The DNA fragment was ligated between the *SphI* and *XbaI* sites of the pHT304.18Z |
| pHT-P<sub>cry2Ab</sub> | A 930-bp DNA fragment containing the promoter of the *cry2Ab* gene was PCR amplified using genomic of the HD1 strain as the template and the primer pair Pcry2Ab-F/ Pcry2Ab-R. The DNA fragment was ligated between the *HindIII* and *BamHI* sites of the pHT304.18Z |
| pHT-P<sub>ami95</sub> | A 199-bp DNA fragment containing the promoter of the *ami* gene was PCR amplified using genomic of the HD1 strain as the template and the primer pair Pamidase-F/ Pamidase-R. The DNA fragment was ligated between the *HindIII* and *BamHI* sites of the pHT304.18Z |
| pHT-P<sub>ami73</sub> | A 343-bp DNA fragment containing the promoter of the *ami* gene was PCR amplified using genomic of the HD73 strain as the template and the primer pair Pamidase2-F/ |
|  | Pamidase-R. The DNA fragment was ligated between the *Hind*III and *Bam*HI sites of the pHT304.18Z |
| pHT-P<sub>*vipR*</sub><sup>-</sup><br>*vipR* | The pHT304.18Z that contains a 2,642-bp DNA fragment containing the promoter and the coding sequence of *vipR* between the *Bam*HI and *Hind*III sites (25) |
| pHT | The pHT304.18 *E. coli*/Bt shuttle vector conferring resistance to ampicillin in *E. coli* and to erythromycin in Bt (52) |
| pHT-*vipR* | The 2,642-bp DNA fragment that contains the promoter and the coding sequence of *vipR* between the *Bam*HI and *Hind*III sites of the pHT-P<sub>*vipR*</sub><sup>-</sup>*vipR* was digested with the same enzymes and cloned in the pHT304.18 |
| pMAD | *E. coli*/Gram-positive bacteria shuttle vector that contains a thermosensitive replication origin and conferring resistance to ampicillin in *E. coli* and to erythromycin in Bt. Designed to perform chromosomal genetic modification (48) |
| pMAD-<br>Δ*spo0A* | The upstream and downstream regions of the *spo0A* gene were amplified by PCR using chromosomal DNA from Bt strain HD73<sup>-</sup> as template and primers *spo0A*-up-1/*spo0A*-up-2Bis and *spo0A*-dn-2/ *spo0A*-dn-2Bis, respectively. The two fragments were fused using complementary sequences located in the primers and resulting DNA fragment was cloned between the *Bam*HI and *Xma*I sites of the pMAD vector |
| pMADK | The DNA fragment containing the minimal thermosensitive replication origin of the pE194, the *BgaB* coding sequence and the pBR222 origin of replication of the pMAD was amplified by PCR using the primer pair pMADopt-kana.FOR/pMADopt-kana.REV and the pMAD as the DNA template. The kanamycin resistance cassette was amplified using the *kanaR*.FOR/ *kanaR*.REV primers and genomic DNA of the Bt407 *spo0A::apha3* strain (53) as DNA template. The two fragments were digested using *Bsa*I and ligated |
| pMADK-<br>Δ*spo0A* | The 1004-bp *Bam*HI-*Xma*I DNA fragment containing the upstream and downstream regions of the HD1 *spo0A* gene of the pMAD-Δ*spo0A* plasmid was digested and cloned into the corresponding restriction sites of the pMADK |

**Table 2:** Strains used in this study.

| Strain | Description | Antibiotic resistance |
| --- | --- | --- |
| HD1 | Bt *kurstaki* HD1 strain, belongs to the serotypes 3a, 3b, 3c (54). |  |
| HD73<sup>-</sup> | AcrySTALLiferous Bt *kurstaki* HD73 strain, cured of the plasmid pHT73 carrying the *cry1Ac* gene (27). |  |
| HD73 | Bt *kurstaki* HD73 strain that contains all the plasmids, belongs to the serotypes 3a, 3b, 3c (54). |  |
| HD73<sup>-</sup> (pP<sub>xyl</sub>) | Strain HD73<sup>-</sup> harboring the pP<sub>xyl</sub>, control strain used to measure the transcriptional activity of the VipR targeted genes. | Km<sup>r</sup> |
| HD73<sup>-</sup> (pP<sub>xyl</sub>, pHT-P<sub>cry1I</sub>) | Strain HD73<sup>-</sup> harboring the pP<sub>xyl</sub>, and the pHT-P<sub>cry1I</sub> plasmids, used to measure the transcriptional activity of the *cry1I* promoter. | Km<sup>r</sup>, Em<sup>r</sup> |
| HD73<sup>-</sup> (pP<sub>xyl</sub>, pHT-P<sub>cry2Aa</sub>) | Strain HD73<sup>-</sup> harboring the pP<sub>xyl</sub>, and the pHT-P<sub>cry2Aa</sub> plasmids, used to measure the transcriptional activity of the *cry2Aa* promoter. | Km<sup>r</sup>, Em<sup>r</sup> |
| HD73<sup>-</sup> (pP<sub>xyl</sub>, pHT-P<sub>cry2Ab</sub>) | Strain HD73<sup>-</sup> harboring the pP<sub>xyl</sub>, and the pHT-P<sub>cry2Ab</sub> plasmids, used to measure the transcriptional activity of the *cry2Ab* promoter. | Km<sup>r</sup>, Em<sup>r</sup> |
| HD73<sup>-</sup> (pP<sub>xyl</sub>, pHT-P<sub>ami95</sub>) | Strain HD73<sup>-</sup> harboring the pP<sub>xyl</sub>, and the pHT-P<sub>ami95</sub> plasmids, used to measure the transcriptional activity of the *ami95* promoter. | Km<sup>r</sup>, Em<sup>r</sup> |
| HD73<sup>-</sup> (pP<sub>xyl</sub>, pHT-P<sub>ami73</sub>) | Strain HD73<sup>-</sup> harboring the pP<sub>xyl</sub>, and the pHT-P<sub>ami73</sub>, used to measure the transcriptional activity of the *ami73* promoter. | Km<sup>r</sup>, Em<sup>r</sup> |
| HD73<sup>-</sup> (pP<sub>xyl</sub>-vipR) | Strain HD73<sup>-</sup> expressing *vipR* under the control of the xylose inducible promoter of *xyIA*, used to | Km<sup>r</sup> |
|  | measure the transcriptional activity of the VipR targeted genes (25). |  |
| HD73<sup>-</sup> (pP<sub>xyI</sub>-vipR, pHT-P<sub>cry1I</sub>) | Strain HD73<sup>-</sup> harboring the pP<sub>xyI</sub>-vipR, and the pHT-P<sub>cry1I</sub> plasmids, used to measure the transcriptional activity of the *cry1I* promoter. | Km<sup>r</sup>, Em<sup>r</sup> |
| HD73<sup>-</sup> (pP<sub>xyI</sub>-vipR, pHT-P<sub>cry2Aa</sub>) | Strain HD73<sup>-</sup> harboring the pP<sub>xyI</sub>-vipR, and the pHT-P<sub>cry2Aa</sub> plasmids, used to measure the transcriptional activity of the *cry2Aa* promoter. | Km<sup>r</sup>, Em<sup>r</sup> |
| HD73<sup>-</sup> (pP<sub>xyI</sub>-vipR, pHT-P<sub>cry2Ab</sub>) | Strain HD73<sup>-</sup> harboring the pP<sub>xyI</sub>-vipR, and the pHT-P<sub>cry2Ab</sub> plasmids, used to measure the transcriptional activity of the *cry2Ab* promoter. | Km<sup>r</sup>, Em<sup>r</sup> |
| HD73<sup>-</sup> (pP<sub>xyI</sub>-vipR, pHT-P<sub>ami95</sub>) | Strain HD73<sup>-</sup> harboring the pP<sub>xyI</sub>-vipR, and the pHT-P<sub>ami95</sub> plasmids, used to measure the transcriptional activity of the *ami95* promoter. | Km<sup>r</sup>, Em<sup>r</sup> |
| HD73<sup>-</sup> (pP<sub>xyI</sub>-vipR, pHT-P<sub>ami73</sub>) | Strain HD73<sup>-</sup> harboring the pP<sub>xyI</sub>-vipR, and the pHT-P<sub>ami73</sub> plasmids, used to measure the transcriptional activity of the *ami73* promoter. | Km<sup>r</sup>, Em<sup>r</sup> |
| HD73 (pHT) | Strain HD73 harboring the pHT plasmid, used as a control strain to study *cry1Ac* transcription and the production of pesticidal protein. | Em<sup>r</sup> |
| HD73 (pHT-vipR) | Strain HD73 harboring the pHT-vipR plasmid, constructed to study *cry1Ac* transcription and the production of pesticidal protein. | Em<sup>r</sup> |
| HD73 Δspo0A | A *spo0A* mutant HD73 strain, defective for its ability to produce a spore. | |
| HD73 Δspo0A (pHT) | Strain HD73 Δspo0A harboring the pHT plasmid, constructed to study *cry1Ac* transcription and the production of pesticidal protein. | Em<sup>r</sup> |
| HD73 $\Delta spo0A$ (pHT-*vipR*) | Strain HD73 $\Delta spo0A$ harboring the pHT-*vipR* plasmid, constructed to study *cry1Ac* transcription and the production of pesticidal protein. | Em<sup>r</sup> |
| HD1 $\Delta spo0A$ | A *spo0A* mutant HD1 strain, defective for its ability to produce a spore. | |

Interestingly, when we searched for a VipR box in the naturally *vipR*-less HD73 strain, one occurrence was found upstream from an *ami* gene located on the pHT73 plasmid (*ami73*). Similarly to the HD1 strain, no VipR box was identified in the chromosome of the HD73 strain. To determine whether the expression of this gene depends on VipR, the pHT-P*_ami73_* plasmid (Table 1) was introduced in the HD73^-^ (pP*_xyl_*) and HD73^-^ (pP*_xyl_*-*vipR*) strains (Table 2). HD73^-^ (pP*_xyl_*, pHT-P*_ami73_*) colonies were white when plated on LB plates containing X-gal and xylose (Fig. 1B). In contrast, the HD73^-^ (pP*_xyl_*-*vipR*, pHT-P*_ami73_*) colonies were blue when *vipR* was expressed indicating that the *ami73* gene was expressed in the HD73 strain only after the introduction of the *vipR* gene.

### The *ami73* and *cry1Ac* genes belong to the same transcription unit

A comparison of all the HD1 and HD73 *ami* loci shows that the promoter and the coding sequence of the *ami73* are identical to those of the *ami65* gene and to the promoter of the transposon-interrupted *ami299\** gene on the pBMB299. A few point mutations and a small insertion differentiate the *ami299* and *ami95* sequences (Fig. S1). Besides the conservation of the *ami* sequences, we noticed that the *ami65, ami95, ami299* and *ami73* genes were located upstream from a *cry1A* gene (Table S1). In the strain HD73, the *ami73* and *cry1Ac* genes are separated by a 289-bp intergenic region that lacks a putative rho-independent transcription terminator (ARNold web server). This suggested that the two genes could belong to the same transcription unit. To test this hypothesis, we performed an RT-PCR experiment with the HD73 strain carrying the pHT-*vipR*, allowing the expression of the regulator under its own promoter, and the HD73 (pHT) carrying the empty pHT304.18 (Tables 1 and 2). RNA were collected at T2 from bacteria grown in LB when VipR was shown to be active (25). The primer pair RTamid-cry1A-F and RTamid-cry1A-R (Table 3) was used to detect the transcription of the putative operon (Fig. 2). An amplicon was produced both in the presence and the absence of *vipR* indicating co-transcription of the genes, although at a low level in the absence of *vipR*. The higher intensity of the PCR product produced with RNA extracted from the *vipR*-carrying strain suggests that VipR increases the transcription of the operon. This result also suggested that VipR can activate the expression of *cry1Ac* early in the stationary phase and independently of the sporulation promoters located in the intergenic region upstream from the *cry1Ac* coding sequence. Indeed, SigE and SigK are not yet activated at this point when bacteria are grown in LB (16). The faint VipR-independent transcription of the *ami73-cry1Ac* operon detected by RT-PCR (Fig. 2) but not in the β-galactosidase assay (Fig. 1B) is presumably due to a weak promoter located upstream of the DNA fragment cloned in the pHT-P*ami73* construct used in the transcription assay.

**Fig. 2.**
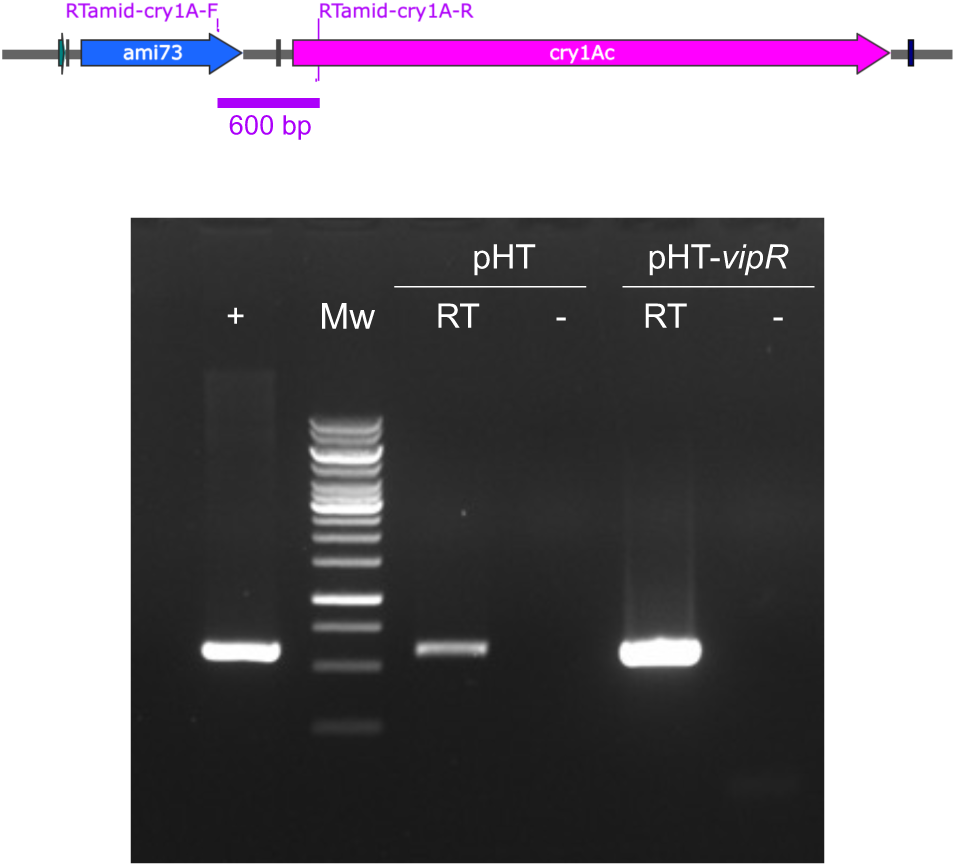
Analysis of the *ami73-cry1Ac* operon in the HD73. A schematic of the *ami73-cry1Ac* locus shows the localisation of the primers used for the RT-PCR experiment. Agarose gel electrophoresis analysis of the production of a 600-bp PCR product obtained with the RTamid-cry1A-F and RTamid-cry1A-R primer pair and HD73 genomic DNA as the DNA template (lanes +), or HD73 (pHT) and HD73 (pHT-vipR) RNA treated (lanes RT), or not (lanes -), with the reverse transcriptase. RNA were collected 2 hours after the onset of stationary phase from bacteria grown in LB medium at 30°C. Mw, GeneRuler^TM^ 1kb DNA molecular weight ladder.

**Table 3:**
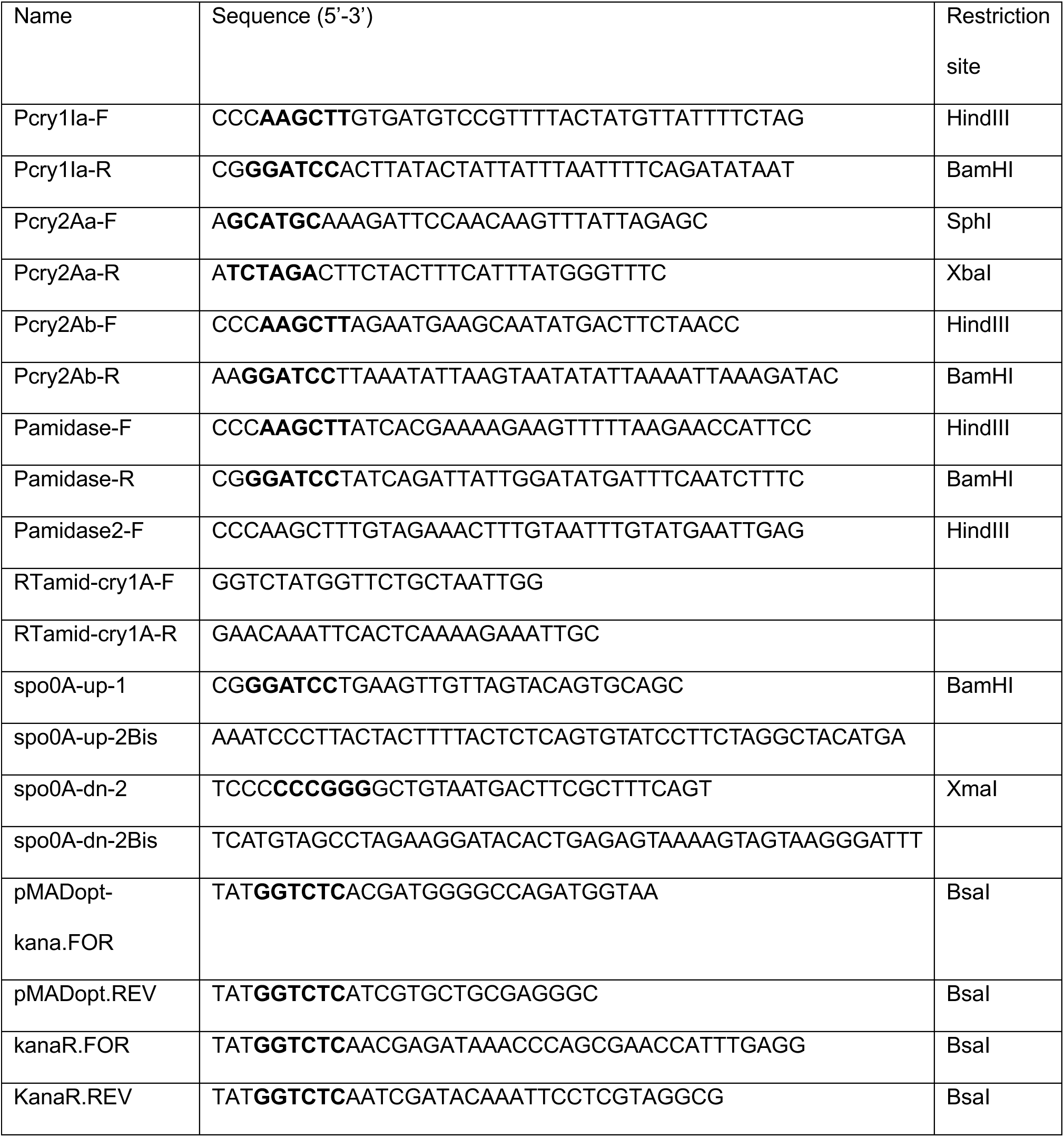
Primers used in this study.

### VipR controls the early production of Cry1Ac

To determine whether the possible VipR-dependent *cry1Ac* expression could lead to the production of the insecticidal toxin in an HD73 strain harboring *vipR*, we analyzed the protein profile of the strains HD73 (pHT-*vipR*) and HD73 (pHT) (Table 2). Bacteria were grown in LB medium and samples were collected from T0 at different time points during the stationary phase. The SDS-PAGE and Western blot analyses indicated that in the presence of *vipR*, a band corresponding to Cry1Ac was present from T0 and accumulated over time (Fig. 3A and 3B). In the absence of *vipR*, Cry1Ac was detected in HD73 (pHT) samples at T11 and later due to SigE- and SigK-dependent *cry1Ac* expression (16). These results indicated that VipR allows the production of the crystal protein from the beginning of the stationary phase of growth. The amount of Cry1Ac protein produced by the two strains was compared by densitometry (Fig. 3C). The quantification of the proportion of the toxin in the 48 h-samples showed that this protein represented, on average, 8,5% of the protein content for the HD73 (pHT) strain. In contrast, this proportion increased to 24,5% for the HD73 (pHT-*vipR*) strain. In agreement with these quantifications, microscopy analysis showed that the *vipR*-carrying strain produced crystals that appeared earlier and seemed bigger than in the *vipR*-less strain (Fig. 3D).

**Fig. 3.**
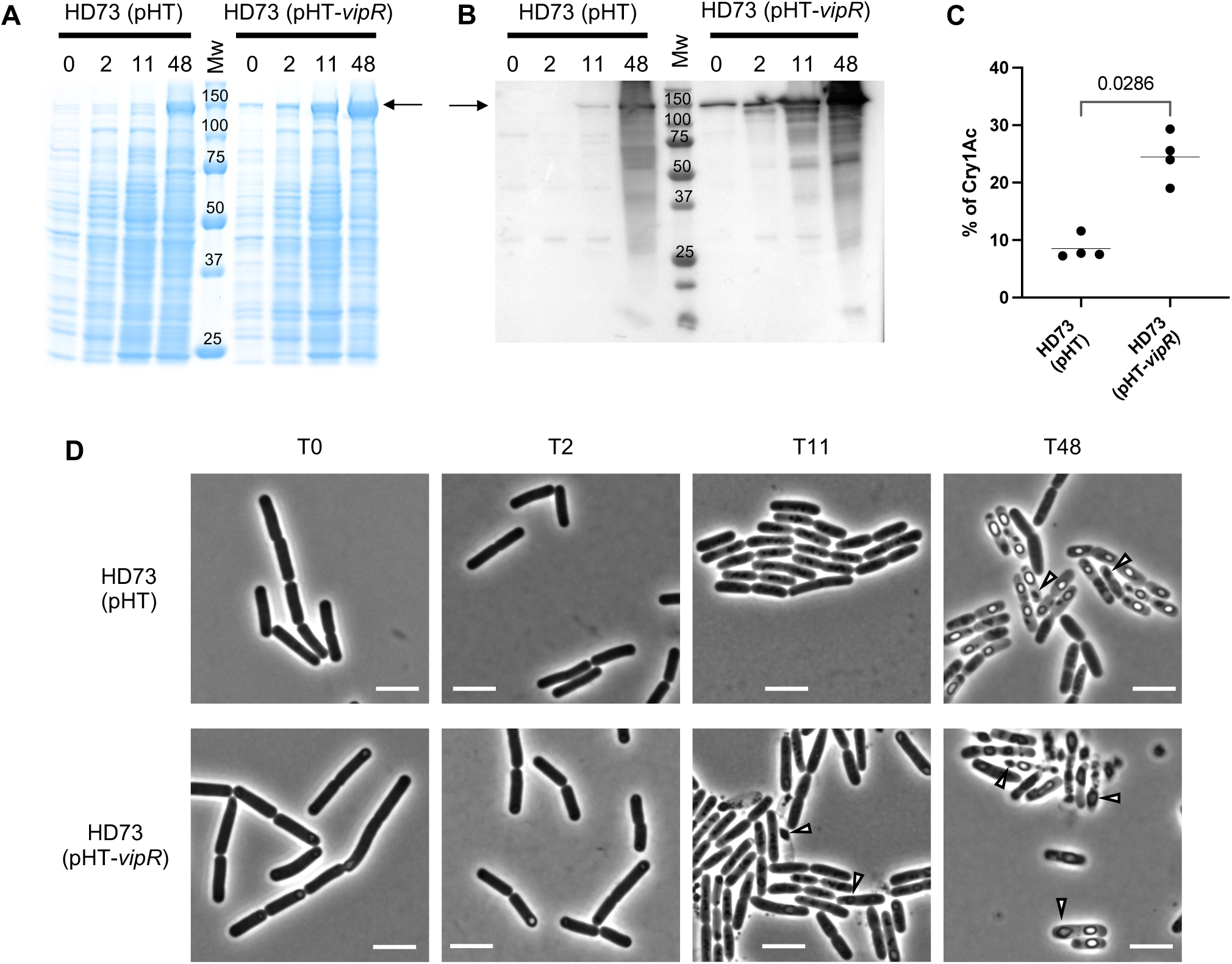
Cry1Ac production in the presence of VipR in the HD73 strain. **A.** SDS-PAGE analysis of total protein extracts of the HD73 (pHT) and HD73 (pHT-*vipR*) strains. Bacteria were grown in LB medium containing 5 µg/mL erythromycin at 30°C and collected at the indicated time (in hours). Proteins were prepared as described in Materials and Methods. 3 µL of sample were loaded in each well. The arrow points to a 133 kDa-migrating protein band that corresponds to Cry1Ac. T0 corresponds to the entry into stationary phase. Mw, protein molecular weight marker. **B.** Western blot analysis of Cry1Ac production by the same strains as in A. 0,3 µL of protein sample was loaded in each well, transferred to PVDF membrane and probed with an antiserum raised against the Bt *aizawai* 7.29 crystal proteins. **C**. Proportion of Cry1Ac in HD73 (pHT) and HD73 (pHT-*vipR*) strains. The proportion of Cry1Ac in SDS-PAGE gel (as in A) was calcultated by densitometry and expressed as the percentage of Cry1Ac in a protein extract at 48h. The data are the result of 4 independent experiments, the bar represents the mean. Result of a Mann-Whitney statistical test is indicated in grey with the P value. **D.** Time course microscopy observations of the HD73 (pHT) and HD73 (pHT-*vipR*) strains grown in LB medium at 30°C for 48h. Arrowheads point to typical Cry1Ac bipyramidal crystals. Scale bar corresponds to 5 µm.

### VipR allows Cry toxin production in a sporulation mutant strain

To assess if Cry1Ac could be produced in a sporulation mutant, we constructed the HD73 Δ*spo0A* strain in which the sporulation process is blocked in its early steps and SigE and SigK are not produced (Table 2) (29, 30) as in *Bacillus subtilis* (31). The protein profile of the strains HD73 Δ*spo0A* (pHT) and HD73 Δ*spo0A* (pHT-*vipR*) (Table 2) was compared when bacteria were grown in LB medium (Fig. 4A and 4B). The HD73 Δ*spo0A* (pHT) did not produce Cry1Ac. In sharp contrast, in the HD73 Δ*spo0A* (pHT-*vipR*) samples, Cry1Ac was detected from the entry into the stationary phase and accumulated over time. Microscopy pictures of the samples showed that the HD73 Δ*spo0A* (pHT) strain did not produce spores or crystals (Fig. 4C). In contrast, crystal-like inclusions were observed in some HD73 Δ*spo0A* (pHT-*vipR*) bacteria in the samples collected at T0 and in more cells over time. At 48 h, large crystals could be observed in the cells. However, the typical bi-pyramidal structure of the crystals produced by the sporulation-deficient cells seemed less conserved than in sporulating bacteria (Fig. 3D and 4C). These results demonstrate that VipR activates the expression and production of the Cry1Ac toxin independently of the sporulation, presumably from the VipR-dependent promoter located upstream from the *ami* gene.

**Fig. 4.**
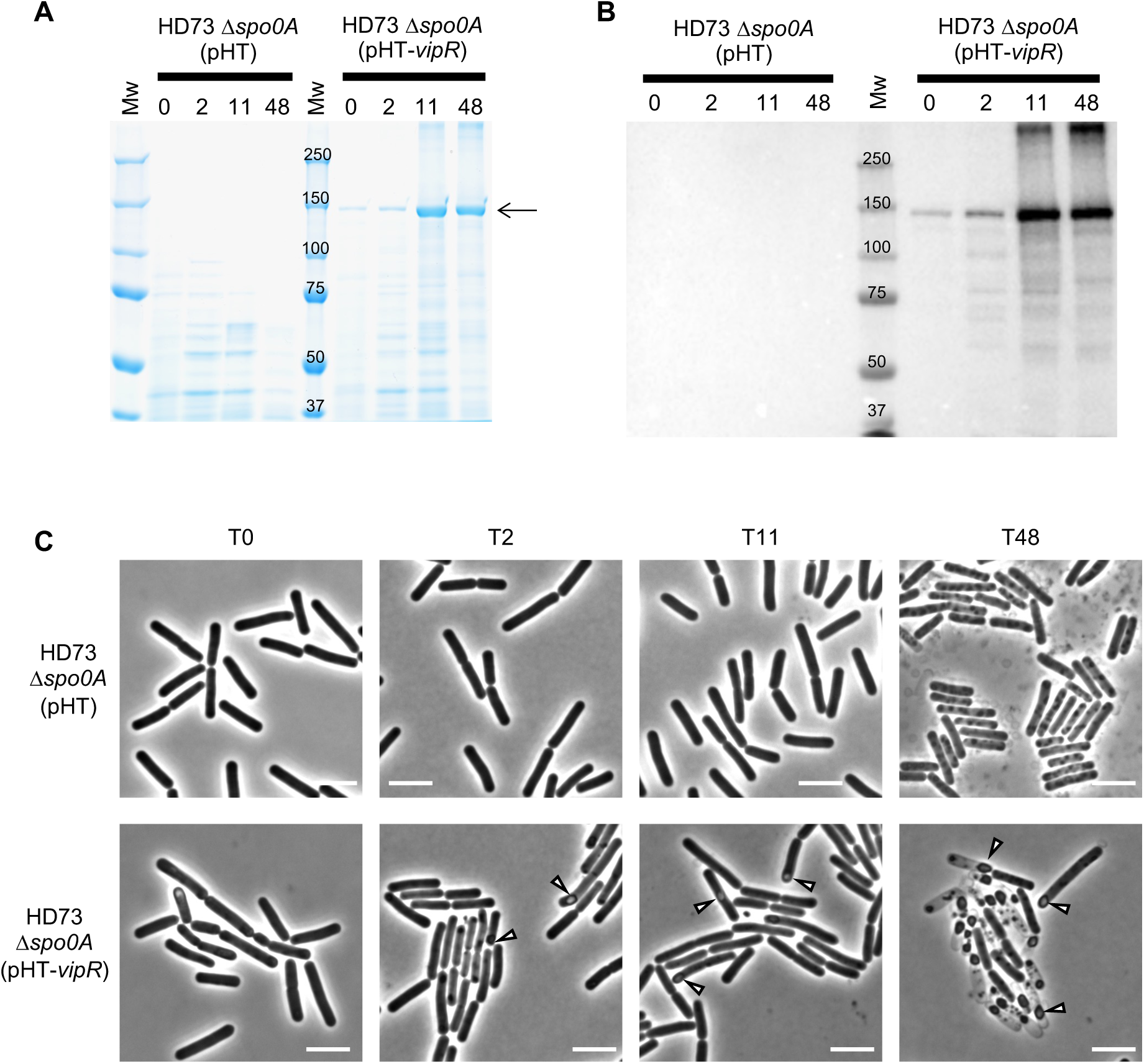
Cry1Ac production in the presence of VipR in the HD73 *Δspo0A* strain. **A.** SDS-PAGE analysis of total protein extracts of the HD73 Δ*spo0A* (pHT) and HD73 Δ*spo0A* (pHT-*vipR*) strains. Bacteria were grown in LB medium containing 5 µg/mL erythromycin at 30°C and collected at the indicated time points (in hours). The arrow points to a 133 kDa-migrating protein band that corresponds to Cry1Ac. Proteins were prepared as described in Materials and Methods. 3 µL of sample were loaded in each well. T0 corresponds to the entry into stationary phase. Mw, protein molecular weight marker. **B.** Western blot analysis of Cry1Ac production by the same strains as in A. 0,3 µL of protein sample was loaded in each well, transferred to PVDF membrane and probed with an antiserum raised against the Bt aizawai 7.29 crystal proteins. **C**. Time course microscopy observations of the HD73 Δ*spo0A* (pHT) and HD73 Δ*spo0A* (pHT-*vipR*) strains grown in LB medium at 30°C for 48h. Arrowheads point to typical Cry1Ac cristalline inclusions. Scale bar corresponds to 5 µm.

### VipR-dependent Cry toxins production in a sporulation mutant HD1 strain

As the work presented above was carried out with our model strain HD73, we wanted to confirm that the VipR-dependent Cry toxin production in Δ*spo0A* bacteria also occurred in the HD1 strain that naturally contains a *vipR* gene. In this strain, VipR might not only control the expression of the three *cry1A* genes but also of *cry1I*, *cry2Ab* and the operon that contains the *cry2Aa* gene (Table S1). Having failed to construct a sporulation mutant in the HD1 strain with the plasmid pMAD, probably because of the selection on erythromycin, as no transformants were obtained, we designed the pMADK plasmid harboring a gene conferring resistance to kanamycin instead of erythromycin (Table 1, Fig. S4 and materials and methods for details). Using this vector, we successfully constructed the HD1 Δ*spo0A* strain (Table 2). While verifying the mutant strain, we noticed that the *cry1Ab* carrying-pBMB65 plasmid had been lost, probably during the culture steps at 40°C to block replication of the thermosensitive pMADK after the recombination events. The HD1 Δ*spo0A* is, therefore, also a Cry1Ab^-^ strain.

Microscopy analyses were conducted to evaluate the production of crystals in the HD1 and HD1 Δ*spo0A* strains (Fig. 5A). The HD1 Δ*spo0A* bacteria did not produce a spore as expected. However, we observed the presence of small inclusions resembling square crystals inside some cells and structures that looked like bipyramidal crystals that were mostly liberated in the growth medium. Large crystals of different shapes were produced as well as spores by the HD1 cells (Fig. 5A). The analysis of the protein profile of the two strains by SDS-PAGE showed that proteins with a molecular weight corresponding to Cry1A and Cry2A (130 and 60 kDa, respectively) were present in both the WT and Δ*spo0A* samples (Fig. 5B), indicating that at least one of the Cry1A and Cry2A toxins was produced by the HD1 Δ*spo0A* strain. The identity of the proteins migrating at 130 kDa and 60 kDa was confirmed by mass spectrometry (Table 4). We also analyzed the two intense 75 kDa protein bands and found a large production of InhA1, InhA2 and InhA3 by the sporulation mutant. A complementary shotgun proteome analysis of the HD1 Δ*spo0A* sample confirmed the production of Cry1Aa, Cry1Ac, Cry1Ia, Cry2Aa, Cry2Ab and Vip3Aa (Table 4). These analyses demonstrate that early and significant production of insecticidal proteins occurs independently of the sporulation in a strain that naturally carries a *vipR* gene and one or several toxin genes that contain a VipR box in their promoter region.

**Fig. 5.**
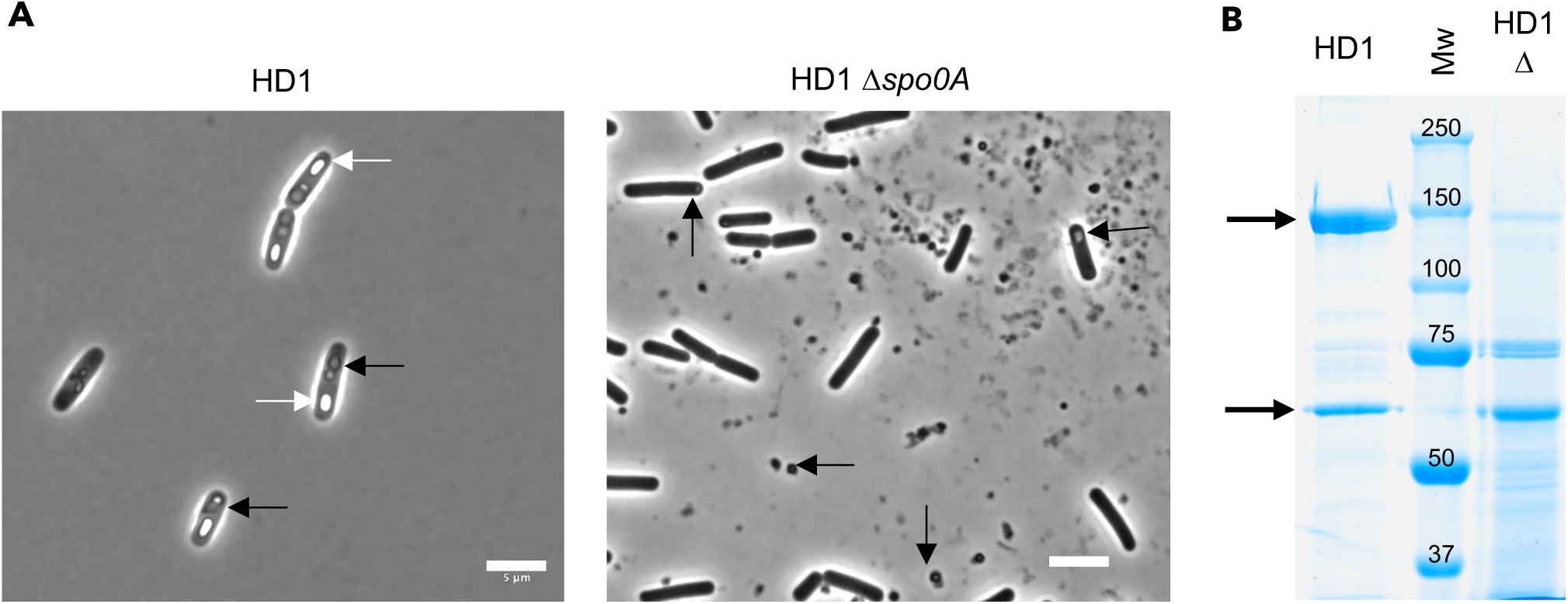
Production of insecticidal proteins by a sporulation-deficient HD1 strain. **A.** Microscopy observations of the HD1 WT and Δspo0A strains grown on LB plates at 30°C for 48 h. Black arrows points to cristalline inclusions, liberated or not. White arrows points to endospores. Scale bar corresponds to 5 µM. **B.** SDS-PAGE analysis of 1,5 µg and 12 µg of a total protein extract of the HD1 (lane HD1) and HD1 Δ*spo0A* (Lane HD1 Δ) strains, respectively. Bacteria were grown on LB medium plates at 30°C for 2 days followed by an incubation at room temperature for 6 days. Arrows point to the 133-kDa Cry1A proteins and the 70-kDa Cry2A proteins. Mw, protein molecular weight marker.

**Table 4:** The most abundant proteins identified in the by LC-MS/MS in the HD1 Δ*spo0A* sample.

| Locus tag | Protein | log(E-value) | Coverage | Spectra | Specific spectra | Unique peptides | Theoretical number of peptides | PAI |
| --- | --- | --- | --- | --- | --- | --- | --- | --- |
| BTK_RS04085 | InhA2 | -494,16 | 48,69 % | 221 | 205 | 116 | 33 | 3,52 |
| BTK_RS08025 | InhA1 | -473,60 | 41,21 % | 213 | 200 | 90 | 30 | 3,00 |
| BTK_RS15040 | InhA3 | -327,05 | 37,23 % | 119 | 110 | 67 | 34 | 1,97 |
|  | Lysozyme C | -481,56 | 89,12 % | 427 | 427 | 106 | 10 | 10,60 |
| BTK_RS32325 | Cry2Ab | -335,79 | 50,55 % | 137 | 99 | 74 | 25 | 2,96 |
| BTK_RS32270 | Cry2Ab | -364,81 | 49,76 % | 135 | 97 | 72 | 26 | 2,77 |
|  | non-ribosomal peptide |  |  |  |  |  |  |  |
| BTK_RS13765 | synthetase | -190,55 | 10,26 % | 63 | 53 | 47 | 214 | 0,22 |
|  | non-ribosomal peptide |  |  |  |  |  |  |  |
| BTK_RS13760 | synthetase | -106,66 | 12,01 % | 37 | 30 | 25 | 86 | 0,29 |
|  | non-ribosomal peptide |  |  |  |  |  |  |  |
| BTK_RS13755 | synthetase | -120,87 | 15,28 % | 32 | 25 | 24 | 60 | 0,40 |
| BTK_RS32295 | Cry1Aa | -241,67 | 25,09 % | 106 | 44 | 47 | 38 | 1,24 |
| BTK_RS31195 | Cry1Ac | -213,94 | 25,57 % | 99 | 40 | 46 | 41 | 1,12 |
| BTK_RS32300 | Cry1Ia | -148,08 | 36,16 % | 57 | 54 | 37 | 26 | 1,42 |
| BTK_RS32255 | Vip3Aa | -217,23 | 34,85 % | 68 | 68 | 49 | 32 | 1,53 |

Coverage, the percentage of the protein sequence covered by the peptides identified; Spectra, the
number of spectra that identified the protein; Specific spectra, the number of specific spectra only
assigned to this protein; Unique peptides, number of unique peptide sequences attributed to this
protein, used to compute the PAI; Theoretical number of tryptic peptides, used in the PAI computation;
Protein Abundance Index (PAI), defines the number of peptides identified divided by the number of
theoretically observable tryptic peptides.

### *In silico* analysis of VipR distribution within the Bt species

While this experimental work has been performed on two Bt *kurstaki* strains, a previous analysis of the presence of the *vipR* gene in Bt strains showed that the regulator gene is present in strains belonging to the *galleriae*, *aizawai*, *chinensis*, *alesti*, *thuringiensis* and *kurstaki*, *finitimus* or *morrisoni* serovars (25). In these strains, a putative VipR box is present upstream from the promoters of the *vipR, vip3, cry1I* and *cry2Ab* genes as well as in the promoter region of the *cry1A* and *cry2Aa* operons when these genes are present. Interestingly, some of these strains carrying *vipR* and VipR boxes may also harbor other insecticidal protein genes not preceded by a VipR box but where typical *sigE* and *sigK* promoters are present. An example of such a strain is the Bt *aizawai* ABTS-1857 (32), where the pCH181-b harbors a *vipR* gene and four insecticidal protein genes, *vip3Aa, cry2Ab, cry1Ia*, and an *ami-cry1Aa* operon with a VipR box in their promoter (Fig. 6 and Table S1), as well as a *cry1Ca* and *cry1Da* genes that do not present a VipR box, but harbor typical SigE and SigK recognition sequences, in their promoters (Table S1 and Fig. S3). This strain also carries another *ami-cry1Ab* operon on the pCH181-e consistent with a VipR-controlled transcription unit as well as a *cry9Ea* gene located on the pCH181-a plasmid and whose promoter seems to be independent of VipR (Table S1 and Fig. S3). Similarly to the *kurstaki* strains, no VipR box could be found in the chromosome of the ABTS-1857 strain.

**Fig. 6.**
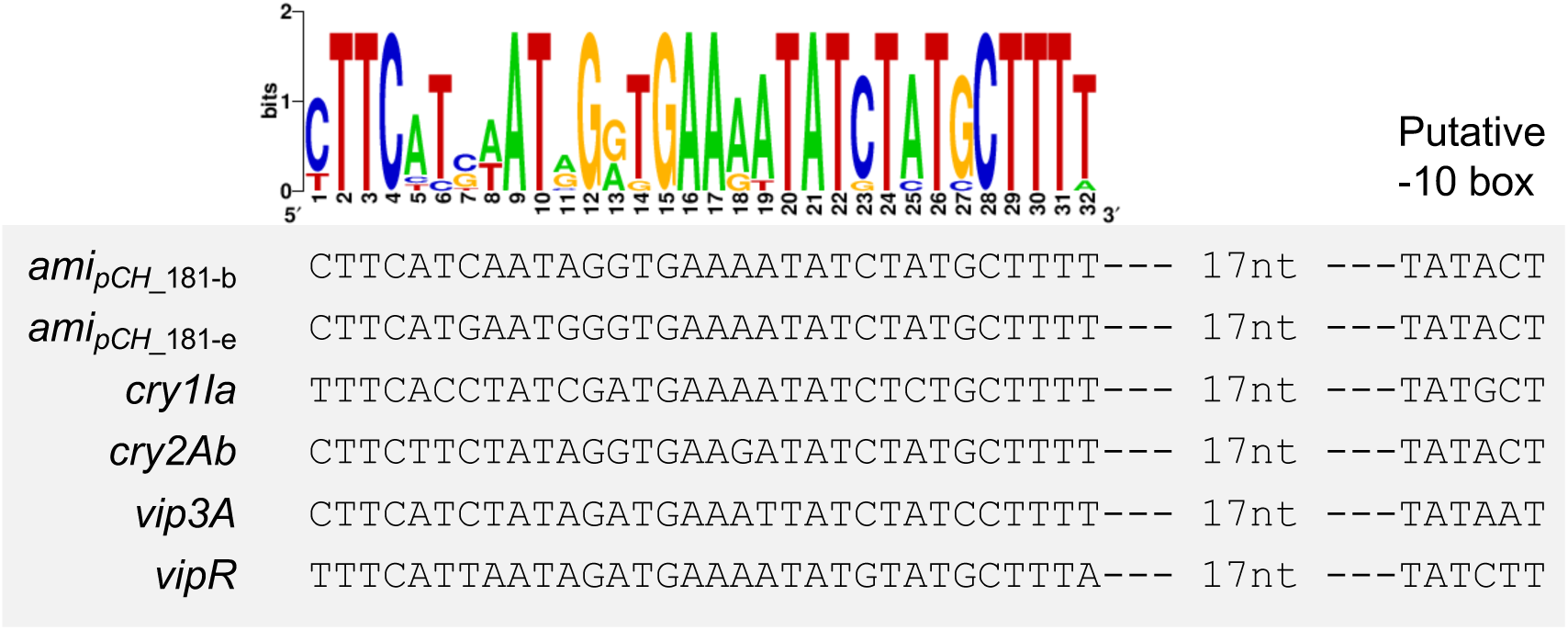
Alignment of the VipR box present in the genome of the Bt *aizawai* ABTS-1857 strain. The name of the gene that contains the conserved sequence in its promoter is indicated on the left. The distance between the putative -10 box and the last nucleotide of the box is indicated. The VipR box web logo constructed with the Bt *kurstaki* HD1 and HD73 sequences is shown on top of the alignment.

## Discussion

For decades, Cry toxin production has been considered to be intimately linked to sporulation, and rightly so. The sporulation sigma factors-dependent *cry* gene expression is a key element enabling the concomitant formation of the spore and the crystal in the mother-cell compartment, with the *cry1* and *cry2* genes being typical examples (9, 12, 13). This finely-tuned gene regulation could be seen as a strategy to produce the perfect set of weapons to successfully achieve the infection of a susceptible insect and the completion of the bacteria life cycle (2). However, identifying the transcription factor CpcR in Bt strain LM1212 (20), the first example of a regulator activating *cry* gene expression in non-sporulating cells, has shed new light on the regulation of these toxin genes. VipR, a transcriptional regulator that belongs to the PCVR family, was previously identified as the transcriptional regulator that controls the expression of the Vip3A insecticidal toxin gene (25). With this work, we demonstrate that VipR is a key player in the expression of insecticidal protein genes (*i.e., cry1A* and *cry2A* genes) whose transcription was previously thought to be exclusively sporulation-dependent (Fig. 1B) (14), as well as the expression of the insecticidal proteins liberated in the growth medium, Vip3A and Cry1I. An unexpected result of our work is related to the expression of the *cry2Ab* gene, considered to date as a cryptic gene (10, 23, 33, 34) eventhough peptides that can be attributed specifically to Cry2Ab have been identified in samples of the DiPel DF (ABTS-351) and XenTari GD (ABTS-1857) commercial products (35). While this protein can be efficiently produced as a P*_cry3A_*-*cry2Ab* promoter-gene fusion (23), two studies have unsuccessfully addressed the natural *cry2Ab* regulation. A first study searched for promoter sequences resembling those preceding other *cry1A* or *cry2Aa* genes but only within the 300 bp upstream from the *cry2Ab* coding sequence (10). No such promoter was identified, which is in agreement with the fact that the VipR-dependent promoter is located more than 700 bp upstream from the *cry2Ab* ORF. The second study used *E. coli* and an HD73 crystal-negative mutant strain as alternative Cry2Ab producers (23). While the whole intergenic region upstream from *cry2Ab*, in which the VipR box is located, was comprised in the DNA construct introduced in these heterologous hosts, Cry2Ab production was not detected. These results are in line with the fact that the *vipR* gene is absent in the genome of these hosts, whereas it is present in strains DiPel DF (ABTS-351) and XenTari GD (ABTS-1857).

An analysis of the VipR-regulated *ami* genes loci drew our attention to an unexpected role of VipR in the expression of the downstream *cry1A* gene, independently of their SigE and SigK-dependent promoters located in the intergenic region between the *ami* and *cry1A* ORFs (Fig. S1). To demonstrate this, we confirmed that the *ami* and the *cry1Ac* genes, previously described in a genomic study as being the minimal *cry1* genetic unit (36) belonged to a same transcription unit (Fig. 2). We also demonstrated that *vipR* enables the expression and production of the endogenous Cry1Ac from the VipR-dependent *ami* promoter (Fig. 3A-B and 4A-B) at the beginning of the stationary phase, hours before the activation of SigE which only occurred approximately 11 h after the onset of stationary phase when bacteria are grown in LB medium (16). A faint transcription of *cry1Ac* originating from the intergenic region upstream of the *cry1A* open reading frame and independent from the sporulation promoters was also described in strain HD73 (16). Spo0A seems to positively regulate this low expression during the stationary phase, but the cause of this activation was not identified. While we also used an HD73 strain in our experiments, we did not observe an early expression and production of Cry1Ac that could originate from the DNA region directly upstream from the *cry1Ac* gene (Fig. 3A-B). We analyzed this DNA sequence for the presence of a VipR box without matching. In line with this idea, the detection of Cry1Ac from T0 was only observed when a VipR expression cassette was introduced in the strain, thus, with a transcriptional activation from the *ami73* promoter.

This report and the studies on the regulator CpcR (14, 20) shed light on a more complex strategy of toxins production in some Bt strains, notably those used as commercial biopesticides. Furthermore, the transcription of VipR and Vip3A is stronger and more precocious in a sporulation mutant, suggesting that the entry into sporulation may be unfavorable for producing the VipR-regulated insecticidal proteins. This phenomenon may also reflect a production of toxins that is triggered by specific environmental signals. Indeed, analysis of the VipR protein sequence shows that the regulator possesses three putative regulatory domains: two phosphoenolpyruvate-sugar phosphotransferase system (PTS) regulation domains (PRD) and one EII-like binding domain (25). These domains are known to be the target of phosphorylation on histidine residues to modulate their activities or their dimerization in response to carbohydrate availability (37). VipR contains histidine residues in its PRD domains that could correspond to the phosphorylated residues in AtxA and Mga (26). Therefore, the activity of VipR might be modulated by environmental signals through the PTS so that the bacteria can produce a large set of toxins to kill their host. This hypothesis would be consistent with the widely accepted chronology of the infection (38). It is demonstrated that the Cry proteins play a major role in the first step of an infection when a susceptible insect feeds and ingests spores and crystals (38, 39). The crystals dissolve in the gut, liberating the toxins leading to toxemia and paralysis of the gut. In some susceptible insects, Cry toxins are sufficient to damage the intestinal epithelium, leading to the host’s death. However, in more resistant insects, the germination of the co-ingested spores is required to produce fatal septicemia. Indeed, spores and the ability to grow after germination in the insect gut have been shown to synergistically increase the insecticidal effect of the Cry proteins (40–42). This effect is due to the production of various virulence factors, mainly belonging to the PlcR regulon, that are produced by the bacterium during the stationary phase (38, 41). The role of Vip3A produced by the HD1 strain during growth has notably been evidenced in increasing significantly the toxicity of the strain towards some lepidopteran insects (4, 40). Indeed, in a S*podoptera exigua* larvae infection assay, the HD1 strain was 16X more toxic than the HD1 Δ*vip3A* strain. However, when streptomycin was added to the insect diet, the toxicity of both strains was reduced, but the toxicity of the HD1 strain was less than 3X more toxic compared to the HD1 Δ*vip3A* strain, demonstrating that the production of Vip3A during bacterial growth in the insect was critical to establish the infection (40). Using our model HD73 strain, we showed that VipR allows the production of Cry1Ac at the beginning of the stationary phase of growth, similarly to the timing of expression of Vip3A. This result suggests that Cry1I, the other pesticidal protein of the HD1 strain released by the bacteria during growth, may also contribute to the pathogenic effect during bacteria growth in the insect larvae, as described for Vip3A.

Similarly, the early VipR-dependent production of the Cry1A and Cry2A proteins (Fig. 5) could be involved in the virulence of the bacteria. It would be interesting to determine the precise contribution of VipR not only in producing the crystal-forming toxins but also in the complete infection cycle of Bt in its ecological niche, the insect larva.

## Materials and methods

### Strains and medium

The Bt *kurstaki* HD73 (43) and HD1 (44) strains belonging to serotype 3 were used in this study. The HD73 crystalliferous strain was designated as HD73, and the HD73 acrystalliferous strain as HD73^−^. *Escherichia coli* strains DH5α or NEB10 were used as the host strain for plasmid construction. *E. coli* strain ET12567 (45) was used to prepare demethylated DNA before transforming Bt by electroporation (46, 47). Bacteria were routinely grown in LB medium at 37°C for *E. coli* and at 30°C for Bt cells. Time 0 was defined as the beginning of the transition phase between the exponential and stationary phases of Bt growth. The following concentrations of antibiotics were used for Bt selection: erythromycin (5 µg/mL) and kanamycin (200 µg/mL). For *E. coli* selection, kanamycin (20 µg/mL) and ampicillin (100 µg/mL) were used. When needed, xylose (20 mM) was added to the culture.

### DNA manipulation and strain construction

The plasmids and bacterial strains used in the study are listed in Tables 1 and 2, respectively. All the DNA manipulation procedures were conducted as described in (25). The primers (Table 3) and sequencing data were provided by Eurofins Genomics.

The pMADK (Table 1), a new version of the pMAD (48), was designed to reduce the size of the original plasmid and change the antibiotic resistance genes used to select the transformation events in *E. coli* and *Bacillus*. Briefly, the pBR222-originating ampicillin resistance gene, the pE194-originating erythromycin resistance gene and *pre* gene, were removed and replaced by a kanamycin resistance cassette that is functional both in *E. coli* and *Bacillus*, yielding the pMADK (Fig. S4). These modifications brought the advantage of increasing the number of unique restriction sites present in the MCS. Plasmid pMADK was used for gene disruption by homologous recombination (48).

### Bioinformatic analyses

The VipR boxes were identified in the plasmid and chromosomal sequences using the following sequence in its IUPAC nucleotide code (https://genome.ucsc.edu/goldenPath/help/iupac.html): (CT)TTC(ATC)(TC)(CGT)(TA)AT(AGC)G(AG)(TG)GAA(AG)(AT)TAT(CG)T(AC)T(CG)CTTT(AT) in the following sequences: the chromosome (CP004069) and pHT73 plasmid (CP004070) of the HD73 strain, in the pBMB95 (CP004875) and pBMB65 (CP004873) plasmids of the HD1 strain and in the chromosome (CP083156), pCH_181-b (CP083158) and pCH_181-e (CP083161) plasmids of the ABTS-1857 strain. When a hit was identified, the presence of a putative -10 box located 17 nucleotides downstream of the last position of the motif was verified to confirm the identification of a VipR box.

The presence of a potential Rho-independent transcription terminator was verified using the ARNold web server (http://rssf.i2bc.paris-saclay.fr/toolbox/arnold/).

### ß-galactosidase assay

The ß-galactosidase activity was monitored using a qualitative method. Cells were streaked on LB plates containing X-gal (50 μg/mL) and xylose 20 mM and incubated at 30°C for 24 h followed by an incubation at 20°C for another 24 h. The blue coloration reflects the activity of the ß-galactosidase.

### mRNA extraction and RT-PCR

RNA was extracted from the Bt HD73 strains grown at 30°C in LB medium and harvested at T2. RNA samples were prepared as described (20) and stored at -80°C before use. The cDNA was generated using random hexamer primers and the Maxima H Minus Reverse transcriptase (Thermo Scientific) on 1 μg of total RNA according to manufacturer instructions (+RT). For each RNA sample, a negative control reaction was performed in the absence of the Reverse transcriptase (-RT). A subsequent PCR was realized with the RTamid-cry1A-F/ RTamid-cry1A-R primer pair on each sample (+RT and -RT) using a standard Taq polymerase (NEB). A PCR positive control was realized on HD73 genomic DNA. The production of an amplicon was visualized on 0,8 % agarose gel in TAE 0,5X buffer.

### SDS-PAGE and Western blot assays

For SDS-PAGE analysis, Bt HD73 strains were grown in LB medium at 30°C in agitated cultures. For each time point, 5 mL culture was collected. Bacteria were separated from the growth medium by centrifugation at 5000 × *g* for 10 min. The bacterial cell pellet was resuspended in 500 µL H_2_O and the suspension was treated with lysozyme (0,1 mg/mL) at 30°C for 30 min. Cells were then disrupted with glass beads in a FastPrep 24 (MP Biomedical). The total cell extract was obtained after the sedimentation of the beads. An equal volume of sample (3 µL for SDS-PAGE and 0,3 µL for Western blot) was separated using SDS-PAGE on a 10% polyacrylamide gel. Gels were either stained with the Readyblue^®^ protein gel stain (Sigma) or subjected to Western blot assays. Total protein extracts of the HD1 strains were produced as follows: strains were grown on LB plates for 2 days at 30°C followed by 6 days at 20°C. Colonies were scrapped from plates, resuspended in 500 µL H_2_O and treated as described above.

Western blot assays were performed as described in (25) using a primary antiserum raised against crystal proteins of the Bt *aizawai* 7.29 strain (49) available in the lab and used at the 1:20 000 dilution.

### Microscopy

Cells were observed with a Zeiss AxioObserver.Z1 inverted fluorescence microscope equipped with a Zeiss AxioCam MRm digital camera. Images were processed with the Zeiss ZEN 2.0 software package.

### LC-MS/MS analysis

5 µg of a total protein extract of the HD1 Δ*spo0A* strain was loaded onto a 10% polyacrylamide gel and the gel was run for a short migration. Proteomics analyses were performed on the PAPPSO platform (http://pappso.inra.fr). The proteins were in-gel-digested with trypsin as described in (50). All peptide samples were analyzed using a timsTOF Pro mass spectrometer coupled with a nanoElute liquid chromatography system (Bruker Daltonik GmbH, Germany). Peptides were directly injected and separated onto an Aurora C18 column (75 μm i.d. × 250 mm, 1.6 μm, 120 Å, Ion Opticks, Australia) in buffer A (0.1% formic acid and 2% acetonitrile). The pep6des were eluted with the following gradient: 2-5% of buffer B (0.1% formic acid in 100% acetonitrile) for 1 min, 5-13% of buffer B for 18 min, 13-19% of buffer B for 7 min, 19-22% of buffer B for 4 min. The column temperature was set to 50 °C. Peptides were introduced into the mass spectrometer via a CaptiveSpray nanoelectrospray ion source (Bruker Daltonik GmbH) at an electrospray voltage of 1.6 kV, an ion source temperature of 180 °C, and a dry gas of 3 L/min. Data collection was carried out using DDA-PASEF mode. The mobility scan range used was 0.7 to 1.10 V·s cm^−2^ (1/ K0), the collisional energy increased linearly from 20.0 eV at 0.60 V·s cm^−2^ (1/K0) to 59.0 eV at 1.60 V·s cm^−2^ (1/K0). Both MS and MS/MS spectra were recorded within a scan range of 100−1700 m/z. TIMS accumulation time was set to 180 ms, with a target precursor intensity of 14,000 arbitrary units (au) and a minimum threshold of 1000 au. Absolute intensity thresholds were set to 10 for mass spectra peak detection and 5000 for mobilogram peak detection. Protein identification was performed by querying MS/MS data against a FASTA format database of the Bt *kurstaki* HD1 (GCF_000717535.1) together with a custom contaminant database using the i2MassChroQ software (version 1.0.19, http://pappso.inrae.fr/). The identified proteins were filtered with a minimum of two different peptides, with a peptide E-value < 10^−2^ and protein E value < 10^−4^.

## Acknowledgments

We are grateful to Alexandra Gruss for sharing information on the pE194 origin of replication for the optimization of the pMAD and Alexandre Bolotin for helpful discussion concerning Bt genome analysis. We thank Christelle Lemy and Samia Ben Rejeb for their technical assistance in constructing the HD73 *spo0A* mutant strain and the PAPPSO facility for the proteomics analysis.

## Figure legends

**Fig. S1-** Schematic of the *ami-cry1A* loci of the Bt HD1 and HD73 strains. The SigE and SigK boxes, and the VipR-regulated promoters are indicated. Putative terminators are shown in white. An alignment of the DNA sequences of the *ami* promoters containing the VipR box is shown at the bottom.

**Fig. S2-** The strains HD73^-^ harboring the plasmids pP_gene_*-lacZ* and the pHT-P*_xyl_*-*vipR* (column « + vipR ») or the pHT-P*_xyl_* (column « - vipR ») were patched on LB medium containing X-gal (50 µg/mL) and xylose 20 mM and grown at 30°C for 24 h. Picture was taken after an additional 24 h of growth at room temperature. The name of the gene whose promoter is fused to *lacZ* is indicated on the left.

**Fig. S3-** Schematic of the *cry9Ea, cry1Ca and cry1Da* loci of the Bt *aizawai* ABTS-1857 strain. The SigE- and SigK-boxes are indicated. Putative terminators are shown in white.

**Fig. S4-** Schematic representing the pMADK and focus on the DNA sequence containing the multiple cloning site (MCS). Unique restriction sites are indicated.

**Table S1:** List of the insecticidal protein genes of the Bt *kurstaki* HD1, *kurstaki* HD73 and *aizawai* ABTS-1857 strains and their plasmid location.

| Strain | Plasmid name | Insecticidal protein transcription units |
| --- | --- | --- |
| *kurstaki* HD1 | pBMB299 | *vip3Aa, cry1Ia, ami299-cry1Aa, orf1-orf2-cry2Aa, cry2Ab* |
|  | pBMB95 | *ami95-cry1Ac* |
|  | pBMB65 | *ami65-cry1Ab* |
| *kurstaki* HD73 | pHT73 | *ami73-cry1Ac* |
| *aizawai* ABTS-1857 | pCH_181-a | *cry9Ea* |
|  | pCH_181-b | *vip3Aa, cry1Ia, ami<sub>pCH_181-b</sub>-cry1Aa, cry2Ab, cry1Ca, cry1Da* |
|  | pCH_181-e | *ami<sub>pCH_181-b</sub>-cry1Ab* |

## References

1. Schnepf E, Crickmore N, Rie JV, Lereclus D, Baum J, Feitelson J, Zeigler DR, Dean DH. 1998. Bacillus thuringiensis and its pesticidal crystal proteins. Microbiol Mol Biol Rev 62:775–806.

2. Agaisse H, Lereclus D. 1995. How does Bacillus thuringiensis produce so much insecticidal crystal protein? J Bacteriol 177:6027–6032.

3. Crickmore N, Berry C, Panneerselvam S, Mishra R, Connor TR, Bonning BC. 2021. A structure-based nomenclature for Bacillus thuringiensis and other bacteria-derived pesticidal proteins. Journal of Invertebrate Pathology 186:107438.

4. Estruch JJ, Warren GW, Mullins MA, Nye GJ, Craig JA, Koziel MG. 1996. Vip3A, a novel Bacillus thuringiensis vegetative insecticidal protein with a wide spectrum of activities against lepidopteran insects. Proc Natl Acad Sci USA 93:5389–5394.

5. Wong HC, Schnepf HE, Whiteley HR. 1983. Transcriptional and translational start sites for the Bacillus thuringiensis crystal protein gene. J Biol Chem 258:1960–1967.

6. Ward ES, Ellar DJ. 1986. Bacillus thuringiensis var. israelensis δ-endotoxin. Journal of Molecular Biology 191:1–11.

7. Zhang J. 1998. Bacillus popilliae cry18Aa operon is transcribed by sigmaE and sigmaK forms of RNA polymerase from a single initiation site. Nucleic Acids Research 26:1288–1293.

8. Poncet S, Dervyn E, Klier A, Rapoport G. 1997. SpoOA represses transcription of the cry toxin genes in Bacillus thuringiensis. Microbiology 143:2743–2751.

9. Bravo A, Agaisse H, Salamitou S, Lereclus D. 1996. Analysis of cryIAa expression in sigE and sigK mutants of Bacillus thuringiensis. Mol Gen Genet 250:734–741.

10. Widner WR, Whiteley HR. 1989. Two highly related insecticidal crystal proteins of Bacillus thuringiensis subsp. kurstaki possess different host range specificities. J Bacteriol 171:965–974.

11. Brown KL. 1993. Transcriptional regulation of the Bacillus thuringiensis subsp. thompsoni crystal protein gene operon. J Bacteriol 175:7951–7957.

12. Brown KL, Whiteley HR. 1988. Isolation of a Bacillus thuringiensis RNA polymerase capable of transcribing crystal protein genes. Proc Natl Acad Sci USA 85:4166–4170.

13. Brown KL, Whiteley HR. 1990. Isolation of the second Bacillus thuringiensis RNA polymerase that transcribes from a crystal protein gene promoter. J Bacteriol 172:6682–6688.

14. Deng C, Peng Q, Song F, Lereclus D. 2014. Regulation of cry Gene Expression in Bacillus thuringiensis. Toxins 6:2194–2209.

15. Porcar M, Déleclusse A, Ibarra JE, Juárez-Pérez V. 2014. Early transcription of Bacillus thuringiensis cry genes in strains active on Lepidopteran species and the role of gene content on their expression. Antonie van Leeuwenhoek 105:1007–1015.

16. Yang H, Wang P, Peng Q, Rong R, Liu C, Lereclus D, Zhang J, Song F, Huang D. 2012. Weak Transcription of the *cry1Ac* Gene in Nonsporulating Bacillus thuringiensis Cells. Appl Environ Microbiol 78:6466–6474.

17. Agaisse H, Lereclus D. 1994. Expression in Bacillus subtilis of the Bacillus thuringiensis cryIIIA toxin gene is not dependent on a sporulation-specific sigma factor and is increased in a spo0A mutant. J Bacteriol 176:4734–4741.

18. Malvar T, Baum JA. 1994. Tn5401 disruption of the spo0F gene, identified by direct chromosomal sequencing, results in CryIIIA overproduction in Bacillus thuringiensis. J Bacteriol 176:4750–4753.

19. Salamitou S, Agaisse H, Bravo A, Lereclus D. 1996. Genetic analysis of cryIIIA gene expression in Bacillus thuringiensis. Microbiology 142:2049–2055.

20. Zhang R, Slamti L, Tong L, Verplaetse E, Ma L, Lemy C, Peng Q, Guo S, Zhang J, Song F, Lereclus D. 2020. The stationary phase regulator CpcR activates *cry* gene expression in non-sporulating cells of *Bacillus thuringiensis*. Mol Microbiol 113:740–754.

21. Srinivas G, Vennison SJ, Sudha SN, Balasubramanian P, Sekar V. 1997. Unique regulation of crystal protein production in Bacillus thuringiensis subsp. yunnanensis is mediated by the cry protein-encoding 103-megadalton plasmid. Appl Environ Microbiol 63:2792–2797.

22. Deng C, Slamti L, Raymond B, Liu G, Lemy C, Gominet M, Yang J, Wang H, Peng Q, Zhang J, Lereclus D, Song F. 2015. Division of labour and terminal differentiation in a novel *Bacillus thuringiensis* strain. The ISME Journal 9:286–296.

23. Dankocsik C, Donovan WP, Jany CS. 1990. Activation of a cryptic crystal protein gene of *Bacillus thuringiensis* subspecies *kurstaki* by gene fusion and determination of the crystal protein insecticidal specificity. Molecular Microbiology 4:2087–2094.

24. Kostichka K, Warren GW, Mullins M, Mullins AD, Palekar NV, Craig JA, Koziel MG, Estruch JJ. 1996. Cloning of a cryV-type insecticidal protein gene from Bacillus thuringiensis: the cryV-encoded protein is expressed early in stationary phase. J Bacteriol 178:2141–2144.

25. Chen H, Verplaetse E, Slamti L, Lereclus D. 2022. Expression of the Bacillus thuringiensis *vip3A* Insecticidal Toxin Gene Is Activated at the Onset of Stationary Phase by VipR, an Autoregulated Transcription Factor. Microbiol Spectr 10:e01205–22.

26. Rom JS, Hart MT, McIver KS. 2021. PRD-Containing Virulence Regulators (PCVRs) in Pathogenic Bacteria. Front Cell Infect Microbiol 11:772874.

27. Wilcks A, Jayaswal N, Lereclus D, Andrup L. 1998. Characterization of plasmid pAW63, a second self-transmissible plasmid in Bacillus thuringiensis subsp. kurstaki HD73. Microbiology, 144:1263–1270.

28. Agaisse H, Lereclus D. 1994. Structural and functional analysis of the promoter region involved in full expression of the cryIIIA toxin gene of Bacillus thuringiensis. Molecular Microbiology 13:97–107.

29. Lereclus D, Agaisse H, Gominet M, Chaufaux J. 1995. Overproduction of Encapsulated Insecticidal Crystal Proteins in a Bacillus thuringiensis spoOA Mutant. Nat Biotechnol 13:67–71.

30. Lereclus D, Agaisse H, Grandvalet C, Salamitou S, Gominet M. 2000. Regulation of toxin and virulence gene transcription in *Bacillus thuringiensis*. Int J Med Microbiol 290:295–299.

31. Hilbert DW, Piggot PJ. 2004. Compartmentalization of Gene Expression during *Bacillus subtilis* Spore Formation. Microbiol Mol Biol Rev 68:234–262.

32. European Food Safety Authority (EFSA), Anastassiadou M, Arena M, Auteri D, Brancato A, Bura L, Carrasco Cabrera L, Chaideftou E, Chiusolo A, Crivellente F, De Lentdecker C, Egsmose M, Fait G, Greco L, Ippolito A, Istace F, Jarrah S, Kardassi D, Leuschner R, Lostia A, Lythgo C, Magrans O, Mangas I, Miron I, Molnar T, Padovani L, Parra Morte JM, Pedersen R, Reich H, Santos M, Sharp R, Szentes C, Terron A, Tiramani M, Vagenende B, Villamar-Bouza L. 2020. Peer review of the pesticide risk assessment of the active substance Bacillus thuringiensis ssp. aizawai strain ABTS-1857. EFS2 18.

33. Crickmore N, Wheeler VC, Ellar DJ. 1994. Use of an operon fusion to induce expression and crystallisation of a Bacillus thuringiensis δ-endotoxin encoded by a cryptic gene. Molec Gen Genet 242:365–368.

34. Jain D, Udayasuriyan V, Arulselvi PI, Dev SS, Sangeetha P. 2006. Cloning, Characterization, and Expression of a New cry2Ab Gene From Bacillus thuringiensis Strain 14-1. ABAB 128:185–194.

35. Caballero J, Jiménez-Moreno N, Orera I, Williams T, Fernández AB, Villanueva M, Ferré J, Caballero P, Ancín-Azpilicueta C. 2020. Unraveling the Composition of Insecticidal Crystal Proteins in Bacillus thuringiensis: a Proteomics Approach. Appl Environ Microbiol 86:e00476–20.

36. Fiedoruk K, Daniluk T, Mahillon J, Leszczynska K, Swiecicka I. 2017. Genetic Environment of cry1 Genes Indicates Their Common Origin. Genome Biology and Evolution 9:2265–2275.

37. Deutscher J, Aké FMD, Derkaoui M, Zébré AC, Cao TN, Bouraoui H, Kentache T, Mokhtari A, Milohanic E, Joyet P. 2014. The Bacterial Phosphoenolpyruvate:Carbohydrate Phosphotransferase System: Regulation by Protein Phosphorylation and Phosphorylation-Dependent Protein-Protein Interactions. Microbiol Mol Biol Rev 78:231–256.

38. Raymond B, Johnston PR, Nielsen-LeRoux C, Lereclus D, Crickmore N. 2010. Bacillus thuringiensis: an impotent pathogen? Trends in Microbiology 18:189–194.

39. Bravo A, Likitvivatanavong S, Gill SS, Soberón M. 2011. Bacillus thuringiensis: A story of a successful bioinsecticide. Insect Biochemistry and Molecular Biology 41:423–431.

40. Donovan WP, Donovan JC, Engleman JT. 2001. Gene knockout demonstrates that vip3A contributes to the pathogenesis of Bacillus thuringiensis toward Agrotis ipsilon and Spodoptera exigua. J Invertebr Pathol 78:45–51.

41. Salamitou S, Ramisse F, Brehélin M, Bourguet D, Gilois N, Gominet M, Hernandez E, Lereclus D. 2000. The plcR regulon is involved in the opportunistic properties of Bacillus thuringiensis and Bacillus cereus in mice and insects. Microbiology (Reading, Engl) 146 ( Pt 11):2825–2832.

42. Li RS, Jarrett P, Burges HD. 1987. Importance of spores, crystals, and δ-endotoxins in the pathogenicity of different varieties of Bacillus thuringiensis in Galleria mellonella and Pieris brassicae. Journal of Invertebrate Pathology 50:277–284.

43. Liu G, Song L, Shu C, Wang P, Deng C, Peng Q, Lereclus D, Wang X, Huang D, Zhang J, Song F. 2013. Complete genome sequence of Bacillus thuringiensis subsp. kurstaki strain HD73. Genome Announc 1:e00080–13.

44. Gonzalez J, Carlton BC. 1980. Patterns of plasmid DNA in crystalliferous and acrystalliferous strains of Bacillus thuringiensis. Plasmid 3:92–98.

45. MacNeil DJ, Gewain KM, Ruby CL, Dezeny G, Gibbons PH, MacNeil T. 1992. Analysis of Streptomyces avermitilis genes required for avermectin biosynthesis utilizing a novel integration vector. Gene 111:61–68.

46. Lereclus D, Arantes O, Chaufaux J, Lecadet M. 1989. Transformation and expression of a cloned delta-endotoxin gene in *Bacillus thuringiensis*. FEMS Microbiol Lett 51:211–217.

47. Macaluso A, Mettus AM. 1991. Efficient transformation of Bacillus thuringiensis requires nonmethylated plasmid DNA. J Bacteriol 173:1353–1356.

48. Arnaud M, Chastanet A, Débarbouillé M. 2004. New Vector for Efficient Allelic Replacement in Naturally Nontransformable, Low-GC-Content, Gram-Positive Bacteria. Appl Environ Microbiol 70:6887–6891.

49. Lecadet M-M, Sanchis V, Menou G, Rabot P, Lereclus D, Chaufaux J, Martouret D. 1988. Identification of a δ-Endotoxin Gene Product Specifically Active against *Spodoptera littoralis* Bdv. among Proteolysed Fractions of the Insecticidal Crystals of *Bacillus thuringiensis* subsp. *aizawai* 7.29. Appl Environ Microbiol 54:2689–2698.

50. Mouawad C, Awad MK, Rodrigues-Machado C, Henry C, Sanchis-Borja V, El Chamy L. 2025. High-Throughput Analysis of the Flagella FliK-Dependent Surfaceome and Secretome in Bacillus thuringiensis. Biology 14:525.

51. Perchat S, Dubois T, Zouhir S, Gominet M, Poncet S, Lemy C, Aumont-Nicaise M, Deutscher J, Gohar M, Nessler S, Lereclus D. 2011. A cell–cell communication system regulates protease production during sporulation in bacteria of the Bacillus cereus group. Molecular Microbiology 82:619–633.

52. Arantes O, Lereclus D. 1991. Construction of cloning vectors for Bacillus thuringiensis. Gene 108:115–119.

53. Sanchis V, Agaisse H, Chaufaux J, Lereclus D. 1996. Construction of new insecticidal Bacillus thuringiensis recombinant strains by using the sporulation non-dependent expression system of cryIIIA and a site specific recombination vector. Journal of Biotechnology 48:81–96.

54. Krywienczyk J, Dulmage HT, Fast PG. 1978. Occurrence of two serologically distinct groups within Bacillus thuringiensis serotype 3 ab var. kurstaki. Journal of Invertebrate Pathology 31:372–375.

